# Population-specific association of *Clock* gene polymorphism with annual cycle timing in stonechats

**DOI:** 10.1101/2022.01.15.476482

**Authors:** Hannah Justen, Timo Hasselmann, Juan Carlos Illera, Kira Delmore, David Serrano, Heiner Flinks, Masayuki Senzaki, Kazuhiro Kawamura, Barbara Helm, Miriam Liedvogel

## Abstract

Timing is essential for survival and reproduction of organisms across the tree of life. The core circadian clock gene *Clk* has been implicated in annual timing and shows highly conserved sequence homology across vertebrates except for one variable region of poly Glutamine repeats. *Clk* genotype varies in some species with latitude, seasonal timing and migration. However, findings are inconsistent, difficult to disentangle from environmental responses, and biased towards high latitudes. Here we combine field data with a common-garden set up to study associations of *Clk* with latitude, migration and annual-cycle timing within the stonechat species complex with trans-equatorial distribution range. Including 950 individuals from nine populations with diverse migratory strategies. Gene diversity was lowest in resident African and Canary Island populations and increased with latitude, independently of migration distance. Repeat length and annual-cycle timing was linked in a population-specific way. Specifically, equatorial African stonechats showed delayed timing with longer repeat length for all annual-cycle stages. Our data suggest that at low latitudes with constant photoperiod, *Clk* genotype might orchestrate a range of consistent, individual chronotypes. In contrast, the influence of *Clk* on annual-cycle timing at higher latitudes might be mediated by its interactions with genes involved in (circadian) photoperiodic pathways.

## Introduction

Time is an essential dimension of the ecological niche of a species ^1–3^. In consequence, organisms have evolved internal (i.e., endogenous) time-keeping mechanisms to anticipate changes in the environment that recur rhythmically with high precision. One such endogenous mechanism are circadian clocks that enable organisms to anticipate diel (i.e. within a 24 h period) changes ^4,5^. Circadian clocks continue to run under experimental conditions (constant light or darkness), and in natural environments they are entrained by diel changes in light intensity and duration. The circadian core clock in many organisms is well characterised on the molecular level as a set of genes that regulate biochemical oscillations with a period of 24 h (here called “circadian genes” ^5–7^). For many species, for example fish, mammals and birds, we observe consistent variation in rhythmic behaviour on the level of the individual e.g., by being consistently early or late relative to the population ^8^. Such so-called “chronotypes” have been linked to allelic variation in circadian genes, mostly, but not exclusively in humans ^9^. The demonstrated links between behaviour and circadian gene regulators have made variation in rhythms particularly accessible to ecologists and evolutionary biologists ^10^. Evolutionary interest so far has mostly focused on the gene *Clk*, a single copy gene that is constrained from evolving fast ^11^. The translated protein CLOCK is a key element in the positive arm of the circadian transcription-translation feedback loop ^12^. *Clk* is highly conserved among vertebrates throughout most of its sequence, except for a C-terminal exonic region that contains an often-variable poly-glutamine (poly-Q; CAG/CAA) trio-repeat motif. Length variation in this region varies between species, but in many species also shows substantial variation between and also within populations ^13^. This variable poly-Q region has been shown to influence the transcription activating potential of the CLOCK protein in *Drosophila* ^14^ and mouse ^15^, where truncated versions of the *Clk* poly-Q domain reduced activation of several transcription factors.

Circadian clocks, and the molecular mechanisms that control them, are also involved in annual time-keeping, in conjunction with circannual clocks. Their main role is measurement of photoperiod (daylength) ^16,17^, whose predictable change over the year provides an external calendar of increasing amplitude pole wards, while being absent at equatorial regions. Annual changes in photoperiod can trigger key life cycle events, such as reproduction or migration, either directly or by entrainment of the circannual rhythm ^18^. Thus, molecular changes in clock genes may affect annual timing either directly via pleiotropic effects on circadian and circannual rhythms, or via influencing photoperiodic responses ^19,20^. For example, in many organisms the circadian clock defines a photo-sensitive time window during which reproduction can be triggered or terminated ^16,17^. Hence, individuals with genetic disposition for early *versus* late-entraining circadian clocks are expected to respond differently to a given daylength ^21^. Recent evidence supports corresponding diel and annual chronotypes, e.g. migration-linked behaviour in a songbird ^22^. However, photoperiod-dependent links between diel and annual time-keeping can be complex, since daylength increases in spring, but decreases in autumn.

So far, some studies have indeed demonstrated a correlation between *Clk* genotype and annual-cycle traits. For example, poly-Q repeat length correlated with timing of migration in fish ^23,24^ and birds ^25,26^, timing of moult ^27^, as well as latitudinal clines and breeding phenology in some species ^28–30^. However, these results varied between study systems and a consistent pattern is lacking, e.g. no such correlation was detected between poly-Q repeat length and latitudinal clines in other bird and fish species ^31–33^. Furthermore, no correlation was found for timing of breeding ^32^ and migration ^34^ in other bird species, and some species were altogether lacking detectable genetic variation at the poly-Q locus ^35^.

This lack of consistency has raised critical responses towards candidate gene approaches ^2,10,31,36^, especially when evaluating genotype-phenotype association of complex behavioural traits that are generally assumed to be polygenically controlled. Furthermore, circadian genes have broad, highly pleiotropic effects, similar to “house-keeping” genes ^10^, and there is scepticism whether selection for a particular trait (e.g., timing of breeding) would modify genes with such broad functions, rather than acting on more specific physiological pathways ^20,37,38^. In addition, caveats have been raised against investigating single candidate genes that are part of wider pathways, like the *Clk* gene ^36,39^, and genotype-phenotype associations based on polymorphism of these genes. The reasons for the observed inconsistency between studies are so far not clear.

Contradicting results of the role of the *Clk* gene in different species emphasizes that results from these candidate gene studies can’t easily be generalized across taxa ^25^. For different organisms, the role of *Clk* likely differs, for example due to differences in circadian organization, in genetic architecture, in photoperiodic environments, or in demands on time-keeping ^2,8,40,41^. Thus, in some organisms diel and annual timing might correlate, whereas in others separate and independent regulation may be beneficial ^21,22,42^. Furthermore, most studies are correlational, so that genetic contributions to timing are entangled with a host of responses to environmental conditions ^43^.

Here, we address uncertainty over *Clk* genotype-phenotype associations using a widely discussed subject, avian annual-cycle timing, by a combination of experimental approaches and comparison of closely related taxa. In birds, variation in allelic diversity and repeat length has been focally studied in the context of environmental and behavioural seasonality, specifically breeding latitude, annual-cycle timing and migratory distance. With increasing latitude and thus intra-annual amplitude in day length, reproductive seasons are delayed and shortened, moult timing is shifted, and migratory distance generally increases ^44^. Convincing support for these predictions, however is lacking ^45,46^ .

Aiming to shed light on so far inconsistent results, we developed an experimental design that paired biogeographic associations with controlled experiments under laboratory conditions ^38^ . We compared closely related species within a species complex across wide latitudinal breeding ranges and use da common-garden set-up to disentangle inherited timing from environmental responses ^43^.

Specifically, we investigated the relationship between *Clk* poly-Q polymorphism and breeding latitude, migration and annual-cycle timing in a well-studied songbird complex with a trans-equatorial distribution range, the stonechat (*Saxicola* ssp.; Figure 1). Taxonomy of the former species *Saxicola torquata* is undergoing rapid reassessment. Currently, Saxicola is considered a species complex, of which we’re studying populations or subspecies assigned to the species *Saxicola rubicola* (Austria, Ireland, Spain, Germany), *Saxicola torquatus* (Kenya, Tanzania), *Saxicola dacotiae* (Canary Islands), *Saxicola stejnegeri* (Japan), and *Saxicola maurus* (Kazakhstan). Henceforth, we here use the term “population” to refer to any group of individuals belonging to these taxa, except where otherwise stated. We compared *Clk* genotypes of nine populations with breeding ranges spanning 55° of latitude, including resident, short- and long-distance migratory populations from Europe, Asia and equatorial Africa (Table 1; ^47,48^). In addition to their latitudinal variation these populations differ in their annual timing of breeding, moult and migration. Furthermore, stonechats breeding at the equator experience a constant 12:12 h light/dark cycle throughout the year (Figure 1), and thus present a natural scenario that allows to contrast populations coping with annually fixed and seasonally changing daylengths in the same species complex, avoiding cross-species comparative noise. Finally, a subset of four populations was studied under simulated European daylength; specifically: residents (Kenya), partial migrants (Ireland), short-distance (Austria) and long-distance migrants (Kazakhstan). Our annual-cycle perspective ranged from late summer (postjuvenile moult), through autumn migration to spring migration. We investigated *Clk*-related differences in timing with the following key objectives:

1. Investigating patterns in allelic diversity. Assuming that *Clk* genotype affects annual-cycle timing, we expect allelic diversity to vary with latitude and migration behaviour (here quantified as mean migration distance). We test this hypothesis using all nine populations. If individuals within the population differ in timing or in their photoperiodic exposure, we would expect to find high allelic diversity in *Clk*. Evolutionarily, such differences could result from fluctuating selection on timing. For increasing breeding latitude and migration distance, we might expect increased allelic diversity due to fluctuating mortality linked to arrival timing and wintering latitude ^49,50^. In equatorial populations, constant photoperiodic conditions, but inter-annually variable breeding opportunities could either lead to canalisation and favour fixation of an optimised circadian *Clk* genotype, or, conversely, diversified time-keeping and hence, a broad range of genetically determined chronotypes.
2. Investigating associations between repeat length and population-level timing. If *Clk* allele lengths affect annual timing, we expect to detect differences in allele lengths with breeding latitude and migration distance in the nine populations of wild birds. Equatorial populations may again deviate from patterns at higher latitudes because of the absence of photoperiodic change.
3. Investigating associations between *Clk* genotype and individual-level timing of annual cycle traits. We test directly for genotype-phenotype associations using the four captive populations kept under identical photoperiodic conditions. Because *Clk* allele lengths might affect annual timing via photoperiodic time measurement, we examine whether relationships between repeat length and timing reverses between autumn (decreasing daylength), and spring (increasing daylength). Finally, because the role of *Clk* in annual timing might differ between populations, e.g., due to changes in photoperiodism, we test for population-specific relationships.

**Table 1.**
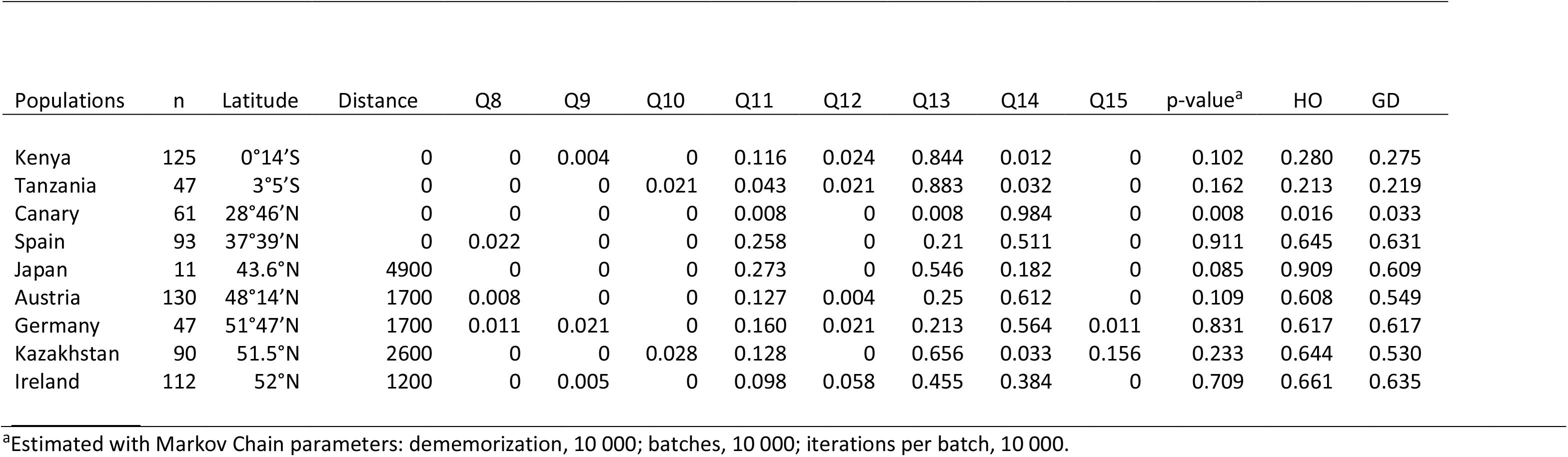
Breeding latitude, migration distance [km], *Clk* poly-Q repeat length frequencies (Q_8-15_), Hardy-Weinberg exact test p-values, observed heterozygosity (HO) and gene diversity (GD) of 9 stonechat (*Saxicola spp*.) populations listed by increasing latitude.

| Populations | n | Latitude | Distance | Q8 | Q9 | Q10 | Q11 | Q12 | Q13 | Q14 | Q15 | p-value<sup>a</sup> | HO | GD |
| --- | --- | --- | --- | --- | --- | --- | --- | --- | --- | --- | --- | --- | --- | --- |
| Kenya | 125 | 0°14'S | 0 | 0 | 0.004 | 0 | 0.116 | 0.024 | 0.844 | 0.012 | 0 | 0.102 | 0.280 | 0.275 |
| Tanzania | 47 | 3°5'S | 0 | 0 | 0 | 0.021 | 0.043 | 0.021 | 0.883 | 0.032 | 0 | 0.162 | 0.213 | 0.219 |
| Canary | 61 | 28°46'N | 0 | 0 | 0 | 0 | 0.008 | 0 | 0.008 | 0.984 | 0 | 0.008 | 0.016 | 0.033 |
| Spain | 93 | 37°39'N | 0 | 0.022 | 0 | 0 | 0.258 | 0 | 0.21 | 0.511 | 0 | 0.911 | 0.645 | 0.631 |
| Japan | 11 | 43.6°N | 4900 | 0 | 0 | 0 | 0.273 | 0 | 0.546 | 0.182 | 0 | 0.085 | 0.909 | 0.609 |
| Austria | 130 | 48°14'N | 1700 | 0.008 | 0 | 0 | 0.127 | 0.004 | 0.25 | 0.612 | 0 | 0.109 | 0.608 | 0.549 |
| Germany | 47 | 51°47'N | 1700 | 0.011 | 0.021 | 0 | 0.160 | 0.021 | 0.213 | 0.564 | 0.011 | 0.831 | 0.617 | 0.617 |
| Kazakhstan | 90 | 51.5°N | 2600 | 0 | 0 | 0.028 | 0.128 | 0 | 0.656 | 0.033 | 0.156 | 0.233 | 0.644 | 0.530 |
| Ireland | 112 | 52°N | 1200 | 0 | 0.005 | 0 | 0.098 | 0.058 | 0.455 | 0.384 | 0 | 0.709 | 0.661 | 0.635 |

<sup>a</sup>Estimated with Markov Chain parameters: dememorization, 10 000; batches, 10 000; iterations per batch, 10 000.

**Figure 1.**
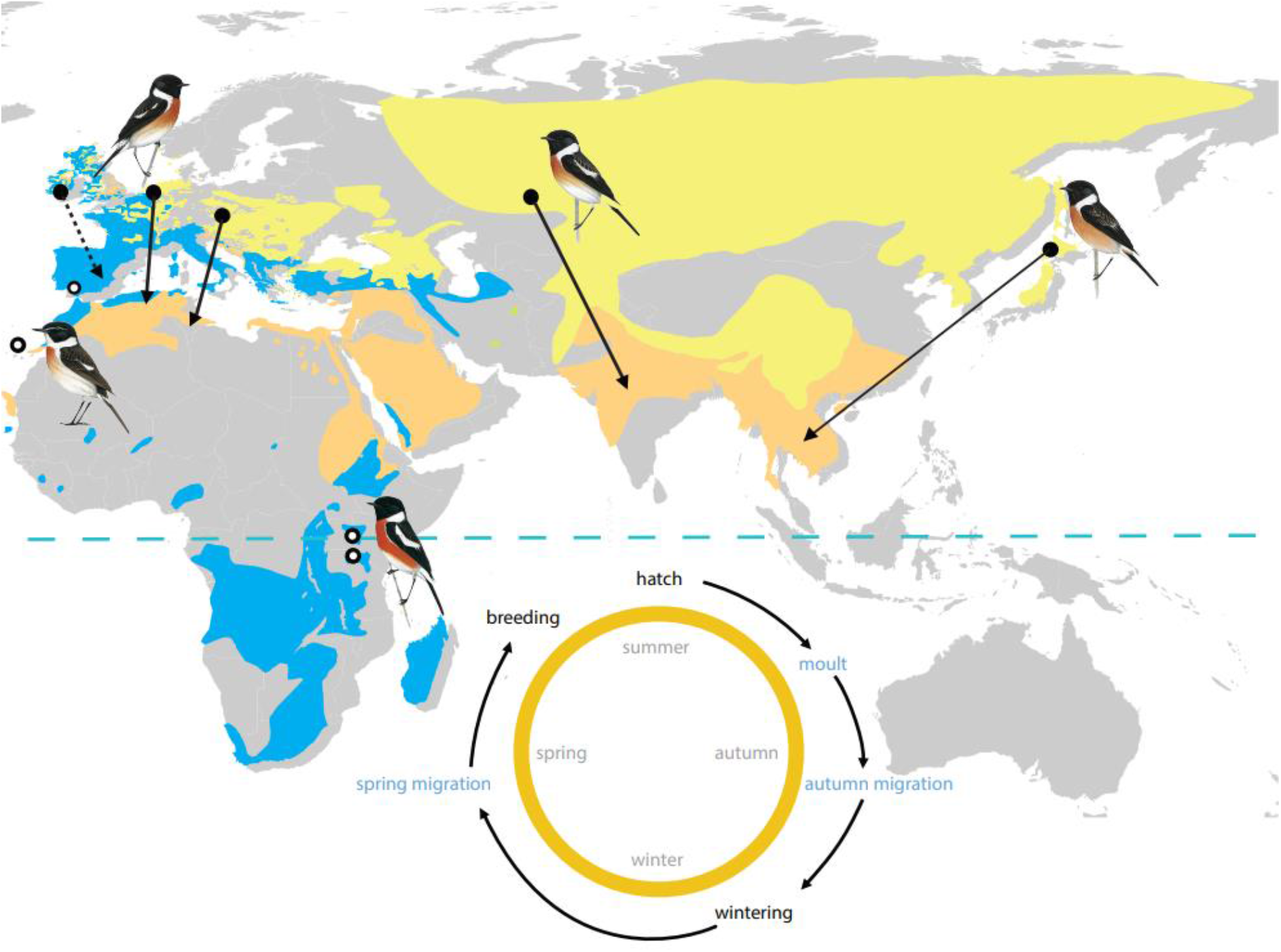
Overview on geographical distribution and migratory behaviour within the *Saxicola* complex. Breeding range (yellow) and wintering range (orange) is shown for migrants, year-round range (blue) is shown for residents; the equator is indicated by a dashed blue line. Breeding location is indicated by filled circles for migratory populations and open circles for residents; arrows depict migratory direction and distance (dotted line indicates partial migrants). The circular inlay schematic illustrates key life history events during the first annual cycle of a stonechat’s life, starting with hatching, followed by moult, autumn migration (in case of migrants), wintering period, return migration in spring, and breeding, before the cycle starts all over again. Focal timing events investigated in this study are highlighted in blue. Bird illustrations show the European *S. rubicola*, the Fuerteventura *S. dacotiae*, the African *S. torquatus*, the Siberian *S. maurus* and the Japanese *S. m. stejnegeri* ^84^ taxa.

## Results

### *Clk* gene diversity, breeding latitude and migration distance

We characterised *Clk* poly-Q allelic lengths variation in nine breeding populations and identified eight length variants ranging from 8 to 15 poly-Q repeats (*Clk* poly-Q_8-15_; subscript indicates the number of poly-Q repeats) found at medium to high frequencies (Table 1). The most common allele (MCA) was Q_13_ for Kenya, Tanzania, Ireland, Japan, and stonechats from Kazakhstan; and Q_14_ for Spain, Austria, Germany and the Canary Islands (Table 1). In both African as well as the Canary Islands populations MCAs accounted for the majority of observed alleles, resulting in reduced population specific allelic diversity: Canary Islands (MCA 98.4%), Kenya (MCA 84.4%) and Tanzania (MCA 88%). In contrast, the contribution of the MCA to overall allelic diversity was considerably lower in stonechats from the European and Asian continent: Kazakhstan (65.6%), Austria (61.2%), Germany (56.4%), Japan (54.5%), Spain (51.1%), Ireland (45.5%) (Table 1). Poly-Q allele frequency within stonechat taxa did not deviate significantly from Hardy-Weinberg equilibrium (p > 0.5; Table 1), except for the Canary Islands population (p < 0.01; Table 1).

*Clk* gene diversity was characterised as observed heterozygosity (defined as frequency of observed number of heterozygotes) as well as within population gene diversity (defined as unbiased gene diversity per sample and locus by Goudet, 2001 ^51^) as well as observed heterozygosity (defined as frequency of observed number of heterozygotes). We analysed *Clk* gene diversity across the geographical range of stonechat populations in the context of breeding latitude and migration distance, two factors that were correlated, albeit not significantly (Pearson’s correlation = 0.56; p = 0.119; df = 7). Equatorial populations from Africa and the Canary Islands showed lower levels of diversity for both measures compared to all other populations; highest diversity levels were found in the Irish population (Table 1). Gene diversity was predicted significantly by breeding latitude, but not migration distance in a joint linear model (ANOVA: latitude: F_2,6_ = 5.417; R^2^ = 0.64; p = 0.017; migration distance: p = 0.694). When we excluded the Canary Island population and restricted our analysis to continental populations, lowest levels of gene diversity were found in the African populations breeding close to the equator (Figure 2).

**Figure 2.**
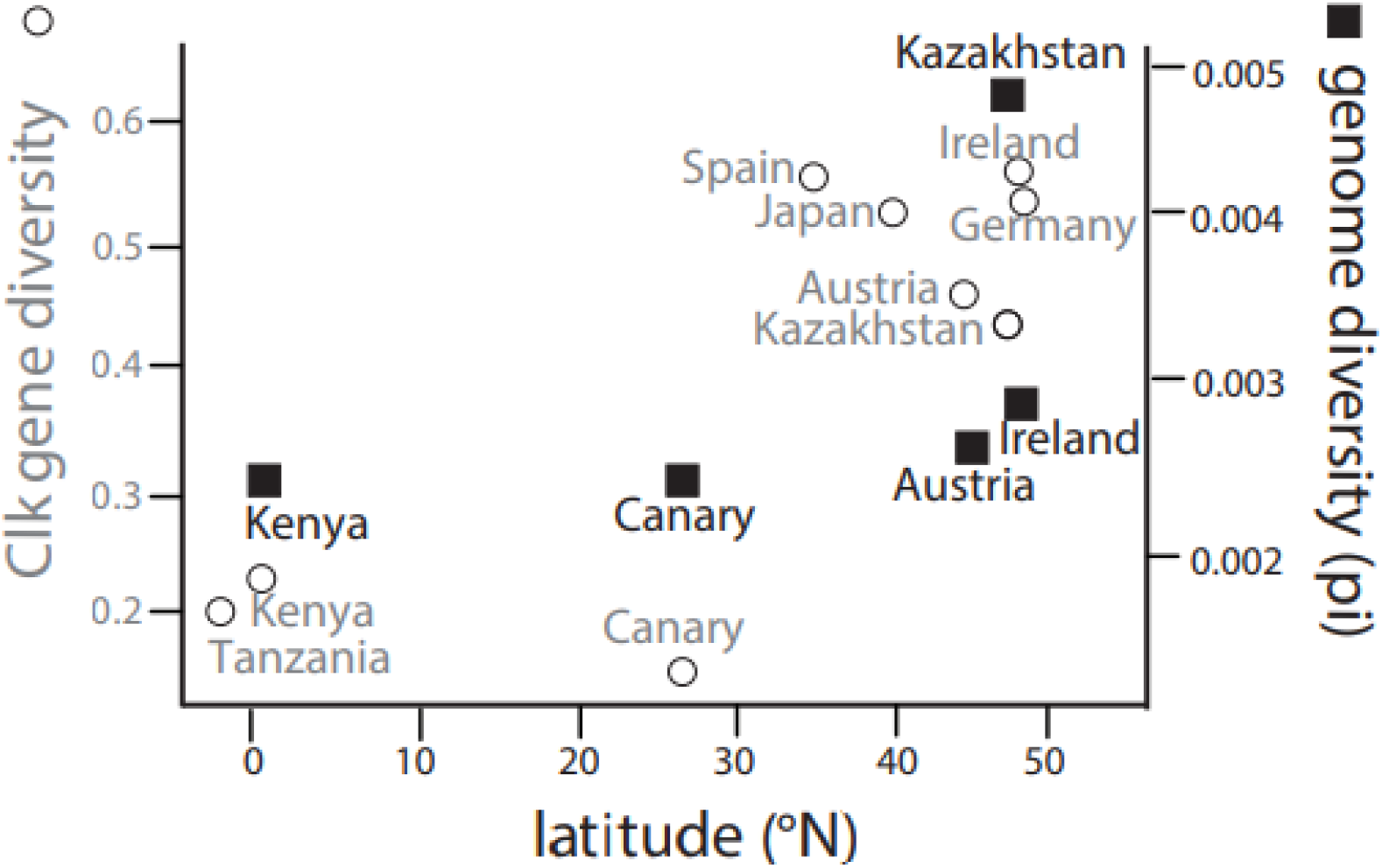
*Clk* locus specific and genome wide diversity of stonechat populations relative to breeding latitude. Gene diversity at the polymorphic *Clk* locus for the nine populations of this study is indicated by open circles (left axis); genome wide nucleotide diversity (autosomal pi) is plotted by black squares (right axis) for five populations with available genome wide statistics ^52^. *Clk* gene diversity is significantly lower in the African populations breeding at the equator than in the populations that breed at higher latitudes in Europe and Asia.

We then tested neutral genome-wide nucleotide diversity (autosomal pi) of five of the included populations (Kenya, Canary Islands, Austria, Ireland and Kazakhstan; estimates from Van Doren et al., 2017) to evaluate if the pattern we observed in *Clk* gene diversity is due to selection in that particular region of the genome or the result of whole genome elevation of nucleotide diversity. We found no correlation between genome-wide nucleotide diversity and breeding latitude nor migration distance (ANOVA: latitude: F_2,2_ = 1.211; R^2^ = 0.55; latitude: p = 0.392; distance: p = 0.380).

### *Clk* repeat length, breeding latitude and migration distance

We detected no significant relationships of breeding latitude or migration distance on *Clk* poly-Q repeat length of individual stonechats (n = 716; linear regression latitude: slope ± SE: 0.38 ± 0.24; df: 6.4; p = 0.161; linear regression migratory distance: slope ± SE: −0.32 ± 0.21; df: 7.24; p = 0.164); see Supplementary Material, Figure S1).

### Genotype-phenotype association: annual-cycle timing

To test for associations between *Clk* genotype, characterised as the mean number of poly-Q repeats at the variable locus, and timing of different focal traits we used the four captive populations and included all available data from a common-garden experiment ^47^ (for sample sizes, see Figure 3).

**Figure 3.**
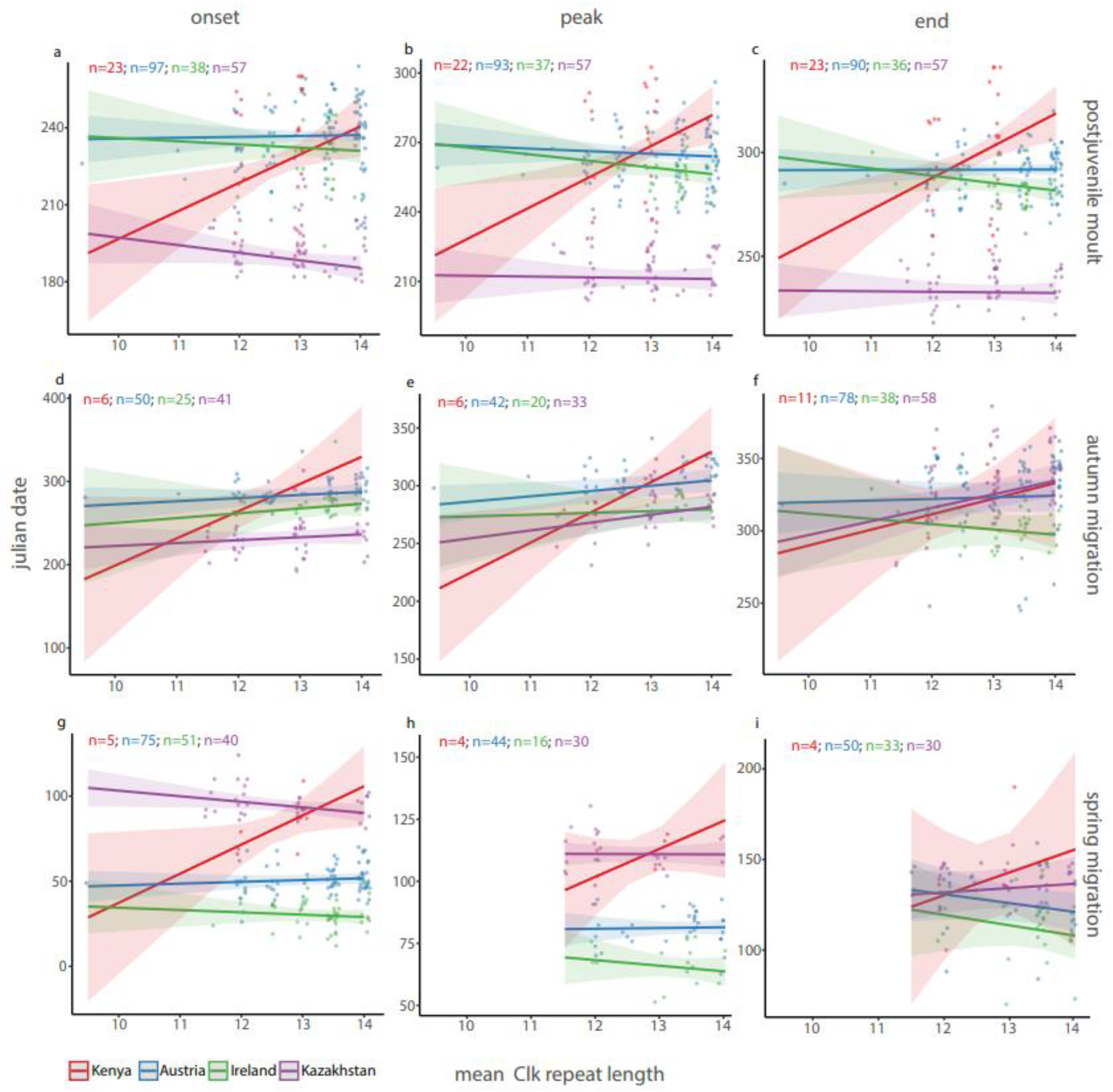
Timing of moult and migration in relation to population-specific *Clk* gene mean repeat length in stonechats. Onset, peak and end of postjuvenile moult (a-c) and of migratory restlessness exhibited during autumn migration (d-f) and spring migration (g-i) are in relation to mean repeat length for four populations: Kenyan stonechat populations plotted in red, Austrian in blue, Irish in green, and Kazakh in purple.

We ran linear mixed effects models including *Clk* repeat length, origin, sex, and hatch date, as well as selected two-way interactions, for onset, peak and end of moult and spring and autumn migratory restlessness.

#### Timing of postjuvenile moult

During this first annual-cycle stage in young birds, timing correlated with *Clk* allele length for onset, peak and end in the Kenyan population (Figure 3a-c; Table 2). These three time points were delayed by 9, 11 and 12 days per additional poly-Q repeat, respectively. In contrast, the relationship between *Clk* mean allele length and timing of the Austrian, Irish and Kazakh populations differed significantly from those of Kenyans and was slightly negative (Figure 3; Table 2). Hatch date had significant effects on moult onset, and this association was population-specific (Table 2). In contrast, sex was not associated moult timing (Table S1).

**Table 2.**
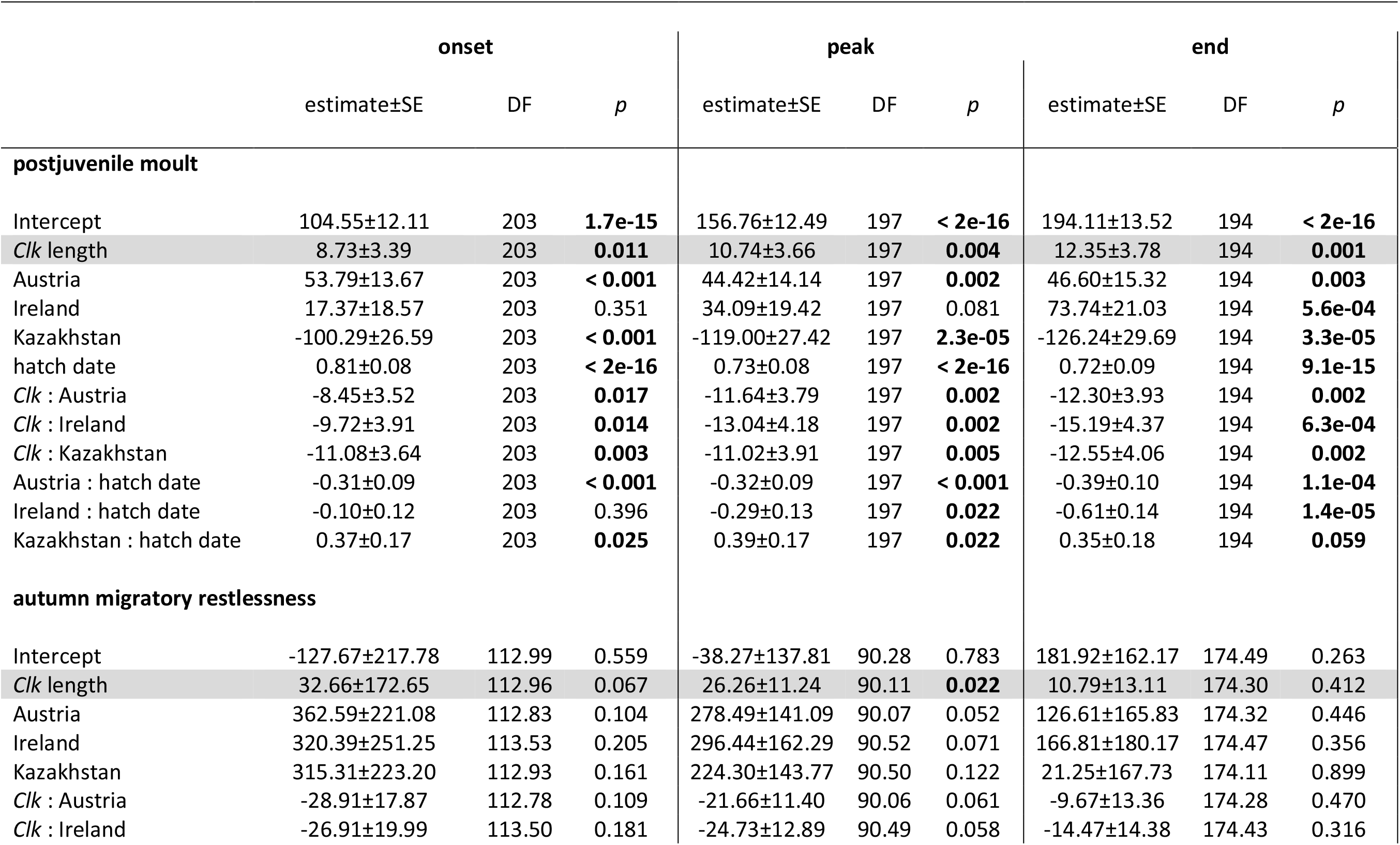
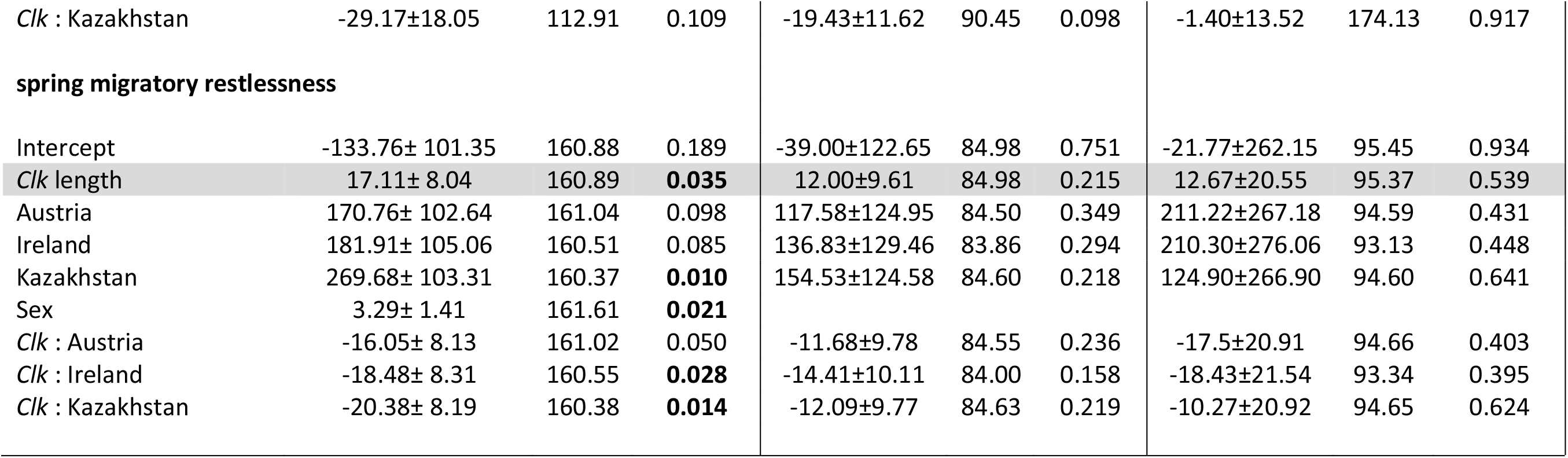
Relationship between *Clk* repeat length and annual cycle timing. Estimates from 9 linear mixed effects models shown for the analysis of postjuvenile moult, and autumn and spring migratory restlessness, based on model selection shown in Supplementary Material Table S1. Reference population for the estimates is Kenyan, reference sex is male.

#### Timing of autumn migratory restlessness

For the subsequent annual-cycle stage, autumn migration, we found a positive relationship between *Clk* allele length and timing (Figure 3d-f; Table 2; Table S1), which did not differ significantly between populations. Slopes were steepest in Kenyan stonechats, whose onset, peak and end were delayed by 33, 25 and 11 days per poly-Q repeat, respectively. Slopes in the remaining populations were far less steep, but overall positive, except for end of autumn restlessness in Irish stonechats (Table 2). Hatching date and sex showed no association with autumn timing.

#### Timing of spring migratory restlessness

Population-specific patterns for spring migratory restlessness were similar (Figure 3g-i; Table 2) to those observed for moult timing. Kenyan stonechats delayed onset, peak and end per poly-Q repeat by 17, 12 and 13 days, respectively, whereas Austrian, Irish and Kazakh populations showed no association or slightly advanced timing with increasing poly-Q repeat. Additionally, we confirmed protandry (i.e. males started migratory restlessness slightly earlier than females) during spring migration (Table 2).

## Discussion

We characterized *Clk* gene poly-Q variation in 950 individuals from nine closely related populations of stonechats, including residents, short-distance and long-distance migrants. The latitudinal range covered by these populations also included equatorial populations, which to our knowledge, were not investigated in previous studies. Our study system thus newly allowed us within the same species-complex to contrast *Clk* gene variation across a latitudinal gradient including the equator.

All stonechat populations were in Hardy-Weinberg equilibrium, suggesting random mating within populations of the same subspecies (p >0.5, Table 1), except for Canary Island stonechats, which diverged from the mainland stonechats 1.6 mya ^53^ and are endemic to the island of Fuerteventura ^54^. A bottleneck event in the Canary Island population after colonization may explain their low genetic diversity ^52^, and their resident behaviour ^55^, along with non-overlapping ranges with other stonechat taxa, may explain the deep genetic differentiation. This history is consistent with significant deviation from Hardy-Weinberg equilibrium (p=0.008; Table 1) and advises caution when interpreting population genetics results from the Canary Island population.

*Clk* gene diversity in different stonechat populations across a latitudinal gradient revealed substantial variation in different *Clk* poly-Q allelic variants ranging from 8 to 15 repeats at the variable Q-locus (Table 1). Variation across study systems varies considerably and in *Saxicola spp.* levels of diversity at the *Clk* locus (here as observed heterozygosity [HO=0.016-0.909]) and frequency of various alleles is comparably high compared to other passerines (e.g. Q_5-8_ in barn swallows [*Hirundo rustica*, HO=0.066] ^30^; Q_10-15_, Q_16_ in bluethroats [*Luscinia svecica*, HO= 0-0.476] ^28^ ; Q_9-14_, Q_15_, Q_16_, Q_17_ in blue tits [*Cyanistes caeruleus*, HO= 0-0.637] ^28^ ; Q_10-15_ in great tits [*Parus major*, HO=0.077] ^35^ ; Q_9-Q13_ in nightingales [*Luscinia megarhynchos*, HO=0.55] ^26^, Q_6-10_ in tree pipit [*Anthus trivialis*], Q_10-13_, Q_15_ in pied flycatcher [*Ficedula hypoleuca*, HO= 0.478] ^26^ ; Q_9_, Q_11-16_ in whinchat [*Saxicola rubetra*, HO=0.125] ^26^). One possible explanation could be that our study comprised populations that varied not only in breeding latitude but also migratory behaviour.

Across stonechats, we found a latitudinal pattern in *Clk* gene diversity, whereby genetic variation was reduced in populations breeding at the equator. This is particularly interesting in a chronobiological context as the scope for light entrainment at the equator dramatically differs from seasonally varying light/dark cycles at higher latitudes. The two equatorial breeding populations from Kenya and Tanzania, which experience constant 12-hour days throughout the year, differed from all other populations included here by significant reduction in *Clk* gene diversity. Previous research on African stonechats has revealed that life-cycle timing shows robust annual cycles, both in free-living birds and under restrictive experimental conditions. Thus, captive African stonechats held under constant 12-hour light-dark cycles retained clear circannual rhythms of breeding condition and moult for several years ^56,57^. Thus, although African stonechats have retained the ability to respond to changing photoperiod ^58^, they appear to rely on an endogenous timing mechanism. Hence, the lower variability in *Clk* repeat length could be signature of selection facilitating adaptation to a constant photoperiod in the birds’ environment resulting in a putatively optimised and less variable genotype ^56,57^.

Circannual rhythms of European stonechat populations that live permanently in the northern hemisphere ^59^ are less rigid, and the substantial changes in photoperiod they experience may play a dominant role for their annual time-keeping ^60,61^. Hence, greater variability in *Clk* repeat length might have resulted from fluctuating inter-annual selection for photoperiodic timing, for example, due to rough winters or sudden weather changes ^49,50^. Thus, we speculate that breeding populations in Germany, Austria, Spain and Ireland require integration of photoperiodism ^20^, and thereby a higher degree of genetic diversity in *Clk* genotype.

We consider it unlikely that the observed genotype-phenotype patterns for the focal *Clk* locus are instead caused by random drift, but specific to this locus. While similar *Clk* gene diversities in Austrian, German and Irish stonechat populations could be due to ongoing gene flow between these populations (e.g. resulting from geographic proximity and breeding dispersal ^52,62^), phylogeny cannot explain the similarly high *Clk* gene diversity we find in Kazakh stonechats, which diverged over 2.5 mya ^53^. Furthermore, the pattern in genetic diversity we observe for the focal *Clk* locus here differs from a genome-wide characterisation across populations investigated here where genome wide patterns of nucleotide diversity show very similar levels across all populations except for Kazakhstan with elevated levels (see Figure 2; ^52^).

Our results on stonechats provide little support for effects of migration on *Clk* gene diversity, independently of breeding latitude. The only indicative of effects of migratory phenotype are the differences in level of heterozygosity between migrants and residents (see HO in Table 1), where all migrant populations show at least 2-fold higher levels of heterozygosity compared to most residents (Table 1, Figure 2). However, reduced levels of *Clk* gene diversity in the population from the Canary Islands is like explained by demographic history and although the Spanish population consists of residents *Clk* gene diversity is comparably high to populations of migrants. In addition, *Clk* gene diversity was significantly predicted only by breeding latitude, but not migration distance.

In a cross-species comparative approach across different trans-Sahara migrants Bazzi et al. (2016) ^46^ hypothesise that selection mechanisms for longer repeats in species with small northern breeding could restrict their postglacially acquired *Clk* diversity, while selection forces in species with larger breeding ranges should be weaker resulting in higher *Clk* diversity. They suggest that analyses of *Clk* gene variation in nonmigratory African relatives of Afro-Palearctic migrants, such as included in our study, should provide additional insights for the hypothesised evolutionary scenario. Results in stonechats, however, are contrary to what Bazzi et al. (2016) hypothesized: genetic diversity in African stonechat is significantly lower compared to migratory stonechat populations, and thus do not provide support for this hypothesis – at least not within the stonechat complex.

A latitudinal cline in *Clk* gene variation with longer repeats at higher latitudes has been demonstrated in various bird systems ^28^, suggesting a functional link between changes in daylength and *Clk* gene variation. Our comparison of mean *Clk* length between populations showed a positive (albeit nonsignificant) trend with longer *Clk* alleles at higher latitudes, confirming earlier studies from different species of birds and fish ^23,28,46^. However, long-distance migratory stonechats from the Kazakh breeding population that breed at similar latitudes as European short-distance migrants, showed high frequencies of the longest *Clk* alleles we observed (15% of alleles are Q_15_, Table 1). This observation points towards contributions of additional factors, such as environmental or climatic variation at the breeding or wintering area, or other characteristics of migratory routes, in correlative studies.

To separate genotype-phenotype associations between *Clk* allele length and annual timing from environmental influences, we compared captive individuals from four populations in a common garden, mimicking European daylength changes. These analyses revealed clear population-specific patterns that depended to lesser degree on time of year. *Clk* allele length in the equatorial stonechats from Kenya correlated positively with timing. Individuals delayed onset, peak and end of the three investigated life-cycle stages, moult and migratory restlessness in autumn and spring, by 9 to 33 days per additional poly-Q repeat. Thus, genotype-phenotype associations were retained across a broad range of photoperiods and in traits as different as seasonal nocturnality and plumage renewal. Although sample sizes varied between populations (range: 4 to 23 individuals), the consistent results suggest that *Clk* allele length is robustly associated with individual chronotypes in the equatorial population. In contrast, the three populations breeding at higher latitudes showed weaker and seasonally variable genotype-phenotype associations. Similar to Kenyan stonechats, high-latitude individuals delayed autumn migratory restlessness with increasing poly-Q repeat numbers, but patterns were inverse for moult, and varied for spring migratory restlessness.

Overall, our results fit well with evidence that circadian clock gene variants can also be relevant for coordination of annual timing, but their roles and effects depend on species, populations and time of year ^63^. More broadly, genetic contributions to annual cycle timing have been shown in several common garden studies, in quantitative genetic analyses and in breeding experiments ^43,60^. Heritability (h^2^) estimates from captive migratory songbirds were medium to high for onset of migratory restlessness in blackcaps *Sylvia atricapilla* (0.34-0.45) and garden warblers *Sylvia borin* (0.67 onset for spring and autumn migration); heritability estimates for termination of migratory activity however were lower (0.16-0.44 in blackcaps ^64,65^; for a summary also see ^66^). Heritability estimates of migration timing traits from the wild are scarce, and estimates of repeatability and heritability are generally moderate, but sometimes lower, for example in Collared flycatchers (*Ficedula albicollis*) ^63^. It is thus clear that to a varying degree, flexibility and non-genetic factors, such as learning, state and ontogenetic factors, also contribute to individual variation ^67^. Inherited timing programs need to integrate information from multiple physiological pathways, such as metabolism and photic input ^68^. Hence, evolutionary change in annual cycle timing can involve several pathways including, but not limited to, genes with known circadian roles such as *Clk* ^69^.

Previous findings from the captive stonechat populations used in this study indicate high heritability estimates of annual timing traits in the *Saxicola* complex ^61,70^. Our new data suggest the possibility that the underlying genetic basis for individual timing might differ between populations from different latitudes. In equatorial populations, variation in annual chronotype might be partly regulated through a limited range of variants in the gene *Clk* that exert major effects, whereas at higher latitudes, *Clk* variants may mainly act in conjunction with photoperiodic pathways, and hence show weaker genotype-phenotype associations. Despite the lack of a mechanistic framework and reservations regarding the candidate genes approach, our results provide important and novel insight into understanding the possible genetic basis of annual-cycle timing.

In conclusion, we found a latitudinal cline in *Clk* gene diversity in a large dataset of 950 individuals distributed over a wide geographical range, confirming findings in other species. Making use of a common garden setting our study highlights that the relationship between *Clk* polymorphism and annual-cycle timing in captive stonechats depended on population and on time of year. Our findings also allow us to speculate that in populations that live under unchanging photoperiods of the equator, *Clk* genotype may be less variable, but exert strong association with annual chronotype. Conversely, at higher latitude, other evolutionary forces may favour *Clk* gene diversity to be higher, but its association with phenotype may be obscured by additional molecular inputs into annual timing that furthermore depend on time of year.

## Material and Methods

### Study populations

The breeding distribution of stonechats covers a wide geographical range (35°S – 75°N; 20°W – 180°E). Different populations exhibit a variety of migratory behaviours (Figure 1) and differ geographically in the timing of annual processes such as breeding, moult and migratory activity ^47,61^. In captivity, differences in timing between free-living populations largely persist when the birds are kept under common-garden captive conditions ^61^, and even resident populations from equatorial Kenya display migratory restlessness, albeit at a low level ^47,61^. Combined, this makes stonechats an ideal system to study associations of *Clk* gene polymorphisms with different processes. Here, we capitalise on the following phenotypically and geographically distinct populations to study *Clk* gene polymorphism within one species complex (Table 1) ^52,53,71^:

We studied two African mainland populations (*Saxicola torquata axillaris* Shelley 1884: 135 individuals originating from Kenya (0°14’S, 36°0’E) and 52 individuals originating from Tanzania (3°5’S, 36°5’E)), which are year-round residents. We further include a resident island population (n=61) endemic on Fuerteventura, Canary Islands, Spain (28°46’N,14°31’W; *Saxicola dacotiae dacotiae* Meade-Waldo 1889; referred to as ‘Canary’ in the main text ^54^).

We further included four populations of European stonechat *Saxicola rubicola ssp*: (i) one population (n=217) sampled in Austria (Neusiedel; 48°14’N,16°22’E; *Saxicola rubicola rubicola* L. 1776), (ii) one population (n=47) from north-west Germany (Borken; 51°47’N,6°01’E *Saxicola rubicola rubicola* L. 1776; ^72^), (iii) one population (n=184) from Ireland (Killarney; 52°N,10°W; *Saxicola rubicola hibernans* Hartert 1910), and (iv) one population (n = 93) from mainland Spain (Seville; 37°39’N, 5°34’W *Saxicola rubicola rubicola* L. 1776). Birds from Austria and Germany are obligatory medium-distance migrants that spend the winter in Mediterranean regions including North Africa. Irish stonechats are partial migrants that either stay on the breeding grounds year round or migrate over short distances ^62,73^. Spanish birds originate from a population of non-migratory stonechats (Serrano, D. unpublished data).

Lastly, we included two populations of long-distance migrants, Eurasian stonechat *Saxicola maurus ssp*: one population (n = 150) from Kazakhstan (Kustanaj; 51.5°N, 63°E; *Saxicola maurus maurus* Pallas 1773) and a second population (n = 11) from Japan (Hokkaido; 43.6°N, 141.23°E; *Saxicola maurus stejnegeri* L. 1766). Kazakh birds migrate to India, southern continental China and North East Africa ^52,74^, while Japanese birds migrate from Hokkaido over a continental path to winter in mainland southeast Asia ^75^.

### Sample origin

Genomic DNA was isolated from blood (n = 925) and feathers (n = 25) using a salt extraction protocol and diluted to a working concentration of 25 ng/μl. We successfully genotyped 950 individuals from the nine study populations for variable *Clk* poly-Q repeat length. Most samples (n = 643), including birds from populations in Austria (n = 217), Ireland (n = 144), Kazakhstan (n = 150) and East Africa (Kenya and Tanzania; n = 132), originated from a common-garden experiment that Eberhard Gwinner initiated in 1981 at the Max Planck Institute for Ornithology in Andechs, Germany ^47,61,76^. Additional genotype samples (n = 307) were collected from birds caught in the wild at their breeding grounds. Specifically, these included populations from the Canary Islands ^54^, Spain (D. Serrano, unpublished data), Germany ^77^ and Japan ^75^.

All methods were carried out in accordance with relevant guidelines and regulations, all experimental procedures were approved and conformed to the relevant regulatory standards under permit (number: 55.2-1 -54-2531-119-05) by the state of Upper Bavaria. The study was carried out in compliance with the ARRIVE guidelines (https://arriveguidelines.org)

### Genetic analyses

Genomic DNA samples were genotyped for length polymorphism in the variable poly-Q repeat region of the *Clk* gene using a polymerase chain reaction (PCR) amplification protocol and lengths characterisation of the variable region. PCR amplification was carried out in a 10μl total volume using a previously published primer set Johnsen *et al.* 2007 ^28^ (forward primer: 5’-labelled with the ‘blue’ fluorescent dye 6-FAM 5’-6-FAM-TGGAGCGGTAATGGTACCAAGTA-3’; reverse primer: 5’- TCAGCTGTGACTGAGCTGGCT-3’). PCR conditions were optimised for stonechats following conditions published in Liedvogel *et al.* 2009 ^29^: Amplification reaction conditions for Taq DNA polymerase catalysed PCR for the stonechats: Mg^2^+= 2 mM; 95°C/2min; 95°C/30s, 56.8°C/30s, 72°C/30s, 40 cycles; 72°C/5min, 4°C hold. PCR products were prepared for capillary electrophoresis by adding HIDI-formadide-LIZ 500 mixture. Length polymorphism of amplified products were analysed using geneious version 10.1.3. Repeat counts of lengths characterised samples was confirmed by Sanger sequencing of 50 samples across populations using an adapted version of a previous published protocol ^28,29^ with following primers: forward: 5’-TTTTCTCAAGGTCAGCAGCTTGT-3’; reverse: 5’- CTGTAGGAACTGTTGCGGGTGCTG-3’). Amplification conditions using Taq DNA polymerase for Sanger sequencing were optimised for the stonechat material: Mg^2^+= 2 mM; 95°C/2min; 95°C/30s, 61.1°C/30s, 74°C/30s, 40 cycles; 74°C/5min, 4°C hold. Nucleotide sequences of with Exo/SAP purified PCR fragment were determined with BigDye Terminator ready reaction mix, version 3.1 (Applied Biosystems) under standard sequencing conditions according to the manufacturer’s protocol ^29^.

### Genotype characterisation and population genetic analyses

We characterised *Clk* genotype as mean allele length (p+q/2; as previously used in comparative studies, ^26,28^) and did additional sensitivity tests using minimum and maximum allele length in our models. Results from these sensitivity tests were qualitatively the same, therefore we only present results from models with mean allele length in the main text. Genotype frequency data for all populations were tested statistically for deviation from Hardy-Weinberg equilibrium using GENEPOP 4.2 (web version; http://genepop.curtin.edu.au/; accessed 1 November 2020). We adapted to use the same Markov chain parameters that were used in previous studies ^28,29^: dememorization (10 000), batches (10 000) and iterations per batch (10 000). For each population we calculated observed heterozygosity as a measure of *Clk* allelic diversity as the proportion of observed number of heterozygotes of the total number of individuals. To account for the differences in sample size we used the program FSTAT version 2.9.3.2 ^51^ to calculate Fsdiv as additional index of unbiased gene diversity.

To examine whether the *Clk* locus is under selection we compared genetic differentiation at the *Clk* locus with presumably neutral, genome wide nucleotide diversity. Levels of genome wide nucleotide diversity were calculated as autosomal nucleotide diversity (pi) from publicly available whole genome re-sequencing data for five of the included stonechat populations (Kenya, Canary Islands, Austria, Ireland and Kazakhstan ^52^). To identify if differences in gene diversity were due to differences in breeding latitude or migratory phenotype, we ran linear models predicting gene diversity by covariables latitude as well as migratory distance.

Within individuals, we tested for a latitudinal effect on *Clk* gene length by running a linear model with *Clk* mean repeat length as response variable, latitude, and migratory distance as predictor variable.

### Phenotypes of captive stonechats

Data on migratory phenotypes were previously presented for our four captive stonechat populations ^47^. Briefly, stonechats were hand-raised in the Max Planck Institute for Ornithology, as offspring of captive or wild stonechats belonging to the Kenyan, Austria, Irish or Kazakh populations (for detail see ^47^). The birds were kept indoors under simulated natural daylength changes in south Germany (47.5° N during the breeding season, and 40° N during winter ^61^). Here we include all birds with available phenotype and genotype data (migration: n=142 for spring and n=188 for autumn; moult: n=215 for onset, n=209 for peak and n=206 for end).

Postjuvenile moult in stonechats is the first moult after hatching and equates to the first prebasic moult ^78^. A precise description of how moult was recorded was published previously ^70^. In brief, immature stonechats were regularly checked for body moult in 19 plumage areas. The number of plumage areas moulting provided a moult score for each date to determine the timing of onset and completion of moult ^70^.

We used nocturnal migratory restlessness behaviour as proxy for migratory timing. Migratory restlessness, the nocturnal activity of captive birds during migration periods, generally mirrors the migratory timing of free-living conspecifics ^79^, although in stonechats and other songbird species, resident populations also show some migratory restlessness ^47^. A detailed description of how migratory restlessness of the stonechats was recorded and characterised is published elsewhere ^47,61^.

### Analysis of Genotype-Phenotype association

To investigate correlation between *Clk* genotype and annual-cycle phenotype we first ran linear mixed effects models in a model selection step. We included all fixed effects [mean *Clk* repeat length, origin, sex, hatch date, as well as two-way interactions of *Clk* repeat length with origin and hatch date], as well as random effects [year, individual] (for full models, see Supplementary Material Table S1). In order to test our hypothesis of population-specific effects of *Clk* genotype, we retained the interaction of *Clk* genotype and origin, but otherwise only included significant predictors in our final mixed-effects model (Table 2). Random effects were always retained.

On the individual level, we tested the genotype effect on timing of postjuvenile moult, spring and autumn migration, as response variables, using three different time-points, i.e. onset, peak and end (given as Julian date). Analysis was performed in RStudio interface of R version 3.3.3. ^80,81^ using packages ‘lme4’ ^82^ and ‘sjPlot’ ^83^.

All trends remained unchanged in alternative models where one individual with a rare short poly-Q repeat length of Q8 was initially excluded. However, Q8 genotype for this individual (as classified by size fragment determination) was confirmed by Sanger sequencing, giving us no reason to exclude it from the analysis.

## Acknowledgements

We thank the Max Planck Society (MPRG grant MFFALIMN0001 to ML) and Eberhard Gwinner, whose previous work on the stonechats and the avian clock forms the basis for this study. Sylvia Kuhn and Andrew Fidler have inspired this work by their piloting investigations. Additional thanks to Benjamin van Doren for helpful discussions concerning the incorporation of the phenotype data, Eva Stukenbrock for comments on an earlier version of this manuscript and all members of the Max Planck Research Group for Behavioural Genomics for their comments and support.

## Author Contributions

BH and ML conceived and planned the study. Molecular work was carried out by HJ and TH, statistical analyses by HJ with input from ML, BH, KD. BH, JCI, DS, HF, MS, KK contributed sample material. HJ, ML and BH wrote the manuscript, all co-authors commented and agreed on the final draft.

Competing Interest Statement: authors declare no competing interests.

