## Supplementary_Material for "Population-specific association of *Clock* gene polymorphism with annual cycle timing in stonechats"

[1] Max Planck Institute for Evolutionary Biology, MPRG Behavioural Genomics, 24306 Plön, Germany; [2] Texas A&M University, 3528 TAMU, College Station, TX, 77843, US; [3] Department of Animal Physiology, University of Cologne, Cologne, Germany; [4] Biodiversity Research Institute (CSIC-Oviedo University-Principality of Asturias), Oviedo University, Spain; [5] Department of Conservation Biology; Estación Biológica de Doñana (EBD-CSIC); Avda Américo Vespucio 26; 41092 Sevilla, Spain; [6] Wagnerstr. 19, D-46354 Suedlohn, Germany; [7] Environmental Earth Science, Hokkaido University, Sapporo, Hokkaido, Japan; [8] Graduate School of Agriculture, Hokkaido University, Kitaku Kita 9 Nishi 9, Sapporo, Hokkaido, Japan; [9] Department of Wildlife Biology, Forestry and Forest Products Research Institute, Matsunosato 1, Tsukuba, Ibaraki 305-8687, Japan; [10] Groningen Institute for Evolutionary Life Sciences, Groningen University, The Netherlands; [11] Institute of Biodiversity, Animal Health and Comparative Medicine, University of Glasgow, UK; [12] Institute of Avian Research 'Vogelwarte Helgoland', 26386 Wilhelmshaven, Germany

**Table S1.** Model selection for assessing the relationship between *Clk* allele length and annual cycle timing. Statistical details from the full nine linear mixed effects models shown for the analysis of postjuvenile moult, autumn and spring migratory restlessness. Reference population for the estimates is Kenyan, reference sex is male.

| | **onset** | | | | **peak** | | | | **end** | | | |
|---|---|---|---|---|---|---|---|---|---|---|---|---|
| | Sum of Squares | Mean of Squares | df | *p* | Sum of Squares | Mean of Squares | df | *p* | Sum of Squares | Mean of Squares | df | *p* |
| **postjuvenile moult** | | | | | | | | | | | | |
| *Clk* length | 3038 | 3038 | 1 | **3.2e-08** | 2428 | 2428.2 | 1 | **1.4e-06** | 2714 | 2714 | 1 | **2.4e-06** |
| Origin | 52531 | 17510 | 3 | **< 2.2e-16** | 75822 | 25274 | 3 | **< 2.2e-16** | 106264 | 35421 | 3 | **< 2.2e-16** |
| hatch date | 32836 | 32836 | 1 | **< 2.2e-16** | 21011 | 21011.4 | 1 | **< 2.2e-16** | 13661 | 13661 | 1 | **< 2.2e-16** |
| Sex | 0 | 0 | 1 | 0.946 | 41 | 41.1 | 1 | 0.517 | 53 | 53 | 1 | 0.498 |
| *Clk* : Origin | 773 | 258 | 3 | **0.041** | 763 | 254.4 | 3 | 0.053 | 995 | 332 | 3 | **0.037** |
| Origin : hatch date | 2718 | 906 | 3 | **4.3e-06** | 2849 | 949.7 | 3 | **5.2e-06** | 4476 | 1492 | 3 | **9.0e-08** |

| | | | | | | | | | | | |
|---|---|---|---|---|---|---|---|---|---|---|---|
| **autumn migratory restlessness** | | | | | | | | | | | |
| *Clk* length | 1.61 | 1.61 | 97.19 | **0.037** | 2.82 | 2.82 | 79.26 | **0.011** | 0.89 | 0.89 | 141.22 | 0.191 |
| Origin | 42.06 | 14.02 | 97.90 | **<2e-16** | 25.28 | 8.76 | 80.07 | **4.3e-10** | 21.66 | 7.22 | 142.24 | **4.9e-08** |
| hatch date | 0.89 | 0.89 | 97.10 | 0.118 | 1.33 | 1.33 | 79.98 | 0.079 | 0.33 | 0.33 | 141.26 | 0.428 |
| Sex | 0.01 | 0.01 | 98.08 | 0.894 | 0.35 | 0.35 | 80.55 | 0.364 | 0.02 | 0.02 | 141.82 | 0.856 |
| *Clk* : Origin | 0.75 | 0.25 | 97.61 | 0.557 | 1.85 | 0.62 | 80.14 | 0.228 | 1.48 | 0.49 | 141.01 | 0.415 |
| **spring migratory restlessness** | | | | | | | | | | | | |
| *Clk* length | 0.31 | 0.31 | 128 | 0.099 | 0.21 | 0.21 | 63.63 | 0.274 | 0.05 | 0.05 | 69.14 | 0.729 |
| Origin | 88.02 | 29.60 | 128 | **2.2e-16** | 40.39 | 13.46 | 59.72 | **<2e-16** | 5.51 | 1.84 | 66.77 | **0.004** |
| hatch date | 0.01 | 0.01 | 128 | **0.008** | 0.07 | 0.07 | 49.79 | 0.524 | 0.13 | 0.13 | 67.78 | 0.558 |
| Sex | 0.81 | 0.81 | 128 | 0.752 | 0.45 | 0.45 | 55.15 | 0.110 | 0.16 | 0.16 | 64.48 | 0.516 |
| *Clk* : Origin | 1.26 | 0.42 | 128 | **0.013** | 0.42 | 0.14 | 48.44 | 0.488 | 0.62 | 0.21 | 62.18 | 0.655 |

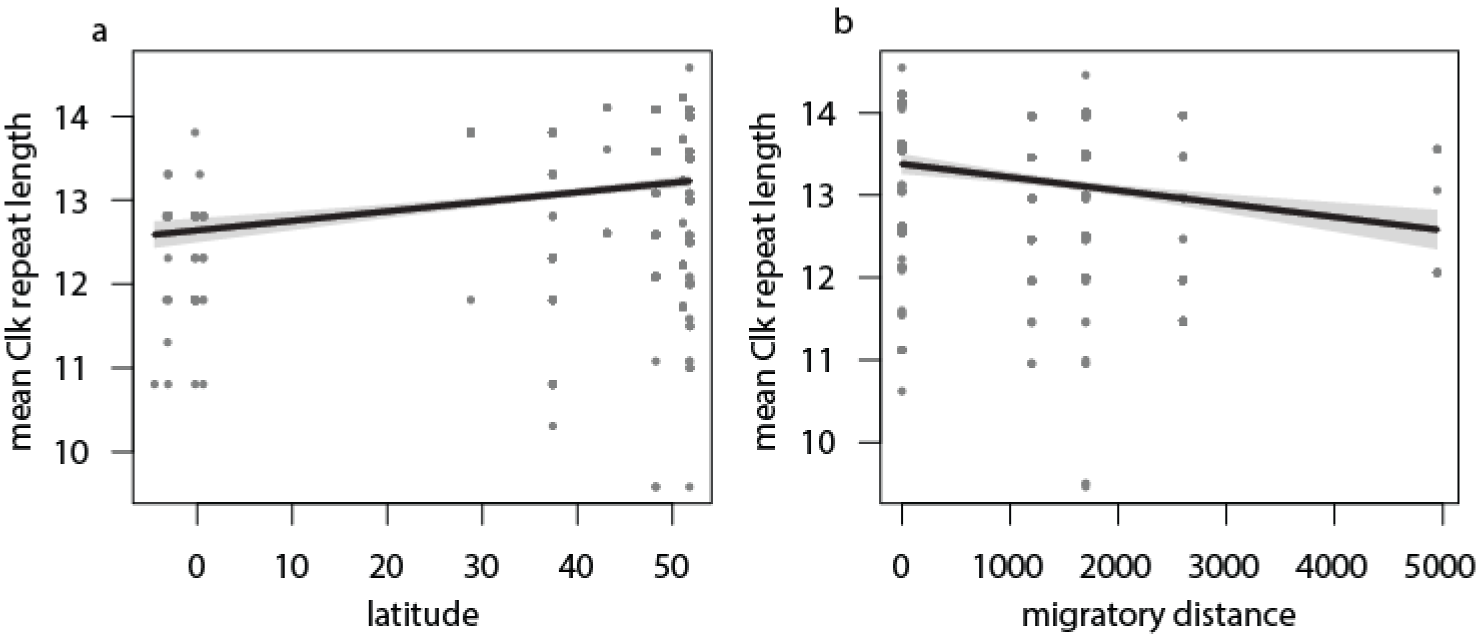

**Figure S1.** Results from linear model investigating breeding latitude (a) and migratory distance (b) of nine stonechat populations in relationship to *Clk* gene mean repeat length. The model shows that mean repeat of individuals increases at higher latitudes and decreases with longer migratory distance ($p < 0.05$).
